# Modeling membrane nanotube morphology: the role of heterogeneity in composition and material properties

**DOI:** 10.1101/373811

**Authors:** Haleh Alimohamadi, Ben Ovryn, Padmini Rangamani

## Abstract

Membrane nanotubes have been identified as dynamic structures for cells to connect over long distances. Nanotubes typically appear as thin and cylindrical tubes, but they may also have a beaded architecture along the tube. In this paper, we study the role of membrane mechanics in governing the architecture of these tubes and show that the formation of bead-like structures along the nanotubes can result from local heterogeneities in the membrane either due to protein aggregation or due to membrane composition. We present numerical results that predict how membrane properties, protein density, and local tension compete to create a phase space that governs the morphology of a nanotube. We also find that there is an energy barrier that prevents two beads from fusing. These results suggest that the membrane-protein interaction, membrane composition, and membrane tension closely govern the tube radius, number of beads, and the bead morphology.

## 1 Introduction

Membrane nanotubes, also referred to tunneling nanotubes, have been identified as intercellular structures that can connect cells over long distances, i.e., hundreds of micrometers (see Fig.1A) [1, 2]. Membrane nanotubes have been observed in a wide variety of cell types [2–5] in addition to the artificial nanotubes that have been produced from lipid vesicles [2, 6]. These nanotubes are typically long and thin cylindrical protrusions with sub-micron diameter and lengths on the order of several hundred microns [1]. In contrast to other types of cellular projections, such as filopodia, which are attached to the substrate and are often function specific [7–9], nanotubes are suspended in the medium and participate in a wide variety of cellular functions [5, 10–13]. Despite increasing observations in the literature highlighting the functional role of membrane nanotubes, the role of membrane mechanics in governing the morphology of these structures has largely remained unexplored.

**Fig. 1:**
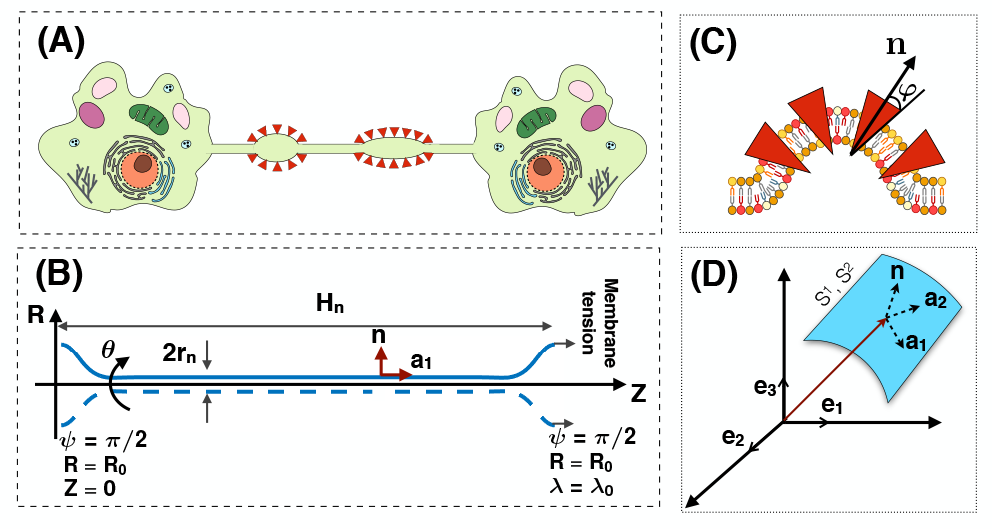
(A) A cartoon showing an intercellular membrane nanotube with local bead-shaped deformations. (B) Axisymmetric coordinates along the membrane nanotube and the boundary conditions used in simulations are shown. (C) A schematic depicting membrane-protein interactions that could lead to the formation of beads along a nanotube. Proteins (shown in red) can aggregate along the membrane to induce local curvature and create a tension heterogeneity, which results in the formation of beads of various sizes and shapes. We assume the dispersed proteins to be cone-shaped such that their meridian makes an angle φ with the normal vector (**n**) to the surface. (D) The coordinate system to define a surface by the tangent basis **a_1_, a_2_** and the normal vector **n**.

Membrane nanotubes have dynamic configurations. Change in the surrounding environment, rearrangement of membrane constituents, or mechanical stresses can result in dramatic shape transformations [1, 14–18]. For instance, in addition to the predominant cylindrical geometry of many membrane nanotubes, the formation of bead-like shapes along membrane nanotubes has been observed in various experiments [12, 19–21]. This morphological transition in nanotubes structures resembles the well-known Rayleigh-Plateau fluid instability, where a change in the surface tension results in the break up of a cylindrical stream of fluid into multiple droplets [22–24]. Similarly, in the case of cylindrical membranes, various studies have shown a tension-driven Rayleigh-type instability, in which a perturbation in the membrane tension leads to the pearling instability [25–30]. Also, the formation of beads along membrane nanotubes could be the result of any lipid bilayer asymmetry such as protein-induced spontaneous curvature, local adsorption of nanoparticles and/or anchored polymers, or induced anisotropic curvatures by membrane inclusions [15, 31–33]. These interactions are accompanied by a change in the local tension of the membrane [34–38]. This suggests that the beading morphology of membrane nanotubes due to external moieties can also be thought of a local version of the tension-driven Rayleigh-type instability.

Recently, a series of elegant modeling studies have proposed the idea that curved proteins and cytoskeletal proteins can induce protrusions along the membrane [39–43]. Separately, experiments are beginning to demonstrate that (a) the composition of a membrane nanotube is not homogeneous [44]; (b) tension due to adhesion of rolling neutrophils can lead to tether formation [44]; and (c) spontaneous curvature along a nanotube can lead to the formation of bead like structures [44–47, 17, 20]. It is not yet clear if such beaded structures are common to membrane nanotubes and if they have a physiological role in cells and tissues. Nonetheless, from a membrane mechanics standpoint, these structures are fascinating to study. Here, we posit that a change in the local membrane tension originating from the heterogeneous distribution of the membrane components can cause the shape transformation from a tubular membrane to a beaded architecture. Therefore, we seek to answer the following questions in this study: How does membrane-protein interaction affect the shape of a nanotube? What is the phase space that governs the morphological transitions of membrane nanotubes? And finally, how do multiple beads interact with each other along the surface of a nanotube? To answer these questions, we used an augmented Helfrich model including the protein density contribution [48–51] with local incompressibility constraints [36, 52] and performed simulations.

## 2 Methods

### 2.1 Assumptions

The plasma membrane is a complex structure; various molecules pack tightly together to form a semi-impermeable shield barrier for the cytoplasm, nucleus, and intercellular organelles [53]. However, under certain assumptions as described below, we can model this complex and heterogeneous surface using a simple mathematical framework to identify some fundamental principles.

- The length scale of the nanotube is assumed to be much larger (~ 20 times) than the thickness of the bilayer. Based on this assumption, we model the membrane as a thin elastic shell [51, 54, 55].
- We assume that the membrane nanotube is at mechanical equilibrium (i.e. no inertia) [56]. Also, because of the high stretching modulus of lipid bilayers [57], we assume that the lipid bilayer is areally incompressible. We use a Lagrange multiplier to implement this constraint [58, 59, 55].
- We assume that the total energy of the system includes the membrane bending energy and a contribution from membrane-protein aggregation in a dilute regime. Thus, the membrane is modeled using an augmented version of Helfrich energy for elastic manifolds including the membrane-protein interaction contributions [48, 60–63]. The aggregation energy of the proteins, which is proportional to the gradient of the protein density, is modeled in our system by a constitutively prescribed hyperbolic tangent function. Also, we assume the lateral phase separation of the proteins is complete, and hence neglect the entropic energy associated with the membrane – protein interactions [64]. A detailed discussion of the full energy can be found in [48, 63, 65].
- We do not explicitly model the diffusion of proteins on the surface and the dynamics of domain formation in this study; rather we assume that diffusion has run its course and we are left with a protein heterogeneity on the surface (see above comment on a hyperbolic tangent). We vary this heterogeneity along the nanotube.
- We assume that the transmembrane proteins are rotationally symmetric. Therefore we ignore the influence of anisotropic membrane inclusions such as BIN-Amphiphysin-Rvs (BAR) domain proteins [41, 40, 66].
- We do not consider the role of forces applied by actin-mediated nanotube formation [40], so that we can focus only on membrane nanotube deformation due to membrane-protein interactions [67–69].
- For simplicity in the numerical simulations, we assume that the membrane in the region of interest is rotationally symmetric and long enough so that boundary effects are ignored (Fig. 1B) [69, 31].

### 2.2 Membrane energy and equations of motion

We model the membrane with two contributions to the strain energy – one from membrane-protein interaction and the other from the membrane bending. The interaction energy of transmembrane proteins with the lipid bilayer is written as a constitutive function of the proein density *σ* (number per unit area). While the exact form of this energy is yet to be experimentally verified, based on thermodynamic arguments, a quadratic dependence of the energy on the local protein density has been proposed as [48, 60, 63, 70–72],

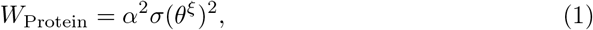

where *W* is the energy per unit area, and *α^2^* is an indicator of the membrane-protein interaction strength 33]. *σ* can depend explicitly on the surface coordinates *θ^ξ^*, where ξ ∈ {1,2} (Fig. 1D) to allow for local heterogeneity. Also, the proteins are assumed to be transmembrane, conical insertions such that the meridian of each protein is at an angle *φ* with the normal vector to the membrane surface (**n**) (Fig. 1C). In the dilute regime, the local induced-spontaneous curvature due to membrane-protein interaction can be modeled as a linar function of the surface protein density as [49, 63]

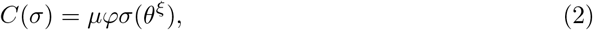

where *μ* is a length scale that represents the lipid-protein specific entropic interactions. Other forms and relationships between the spontaneous curvature and the protein density are explored in [48, 73].

The energy associated with membrane bending due o the isotropic spontaneous curvature is given by the Helfrich Hamiltonian, modified to include the heterogeneous membrane properties as [51, 74, 36]

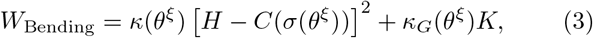

where H is the local mean curvature and K is the local Gaussian curvature. *κ* and *κ_G_* are bending and Gaussian moduli respectively, which in the general case can vary along the surface coordinate *θ^ξ^* [36, 75, 74]. Hence, the total energy of the membrane including both bending and protein contributions is given by

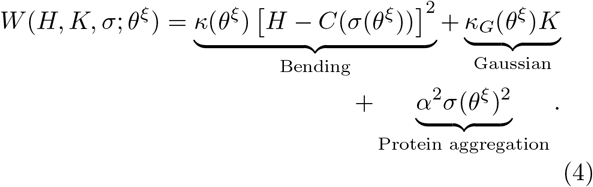

Assuming that the membrane is at mechanical equilibrium at all times, a local balance of forces normal to the membrane, subject to the energy density given in Eq. (4) yields the so-called “shape equation” [49, 50, 76]

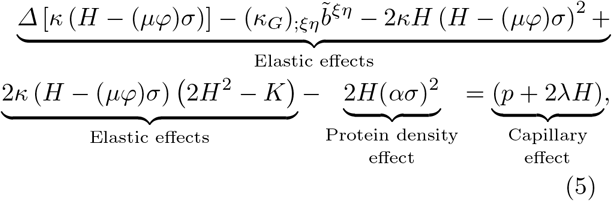

where *Δ* is the surface Laplacian operator, *p* is the pressure difference across the membrane, λ is interpreted as the membrane tension [36, 55], ()_;_ is the covariant derivative with the respect to the surface metric, and 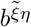 is the co-factor of the curvature tensor.

A local balance of forces tangent to the membrane, which enforces the incompressibility condition in a heterogeneous membrane, yields the spatial variation of membrane tension λ [49, 36, 52, 77]

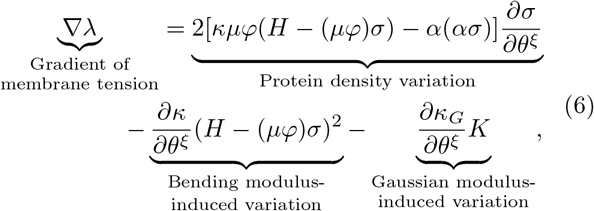

where ∇ is is the partial derivative with respect to the coordinate (*θ^ξ^*). For the sake of brevity, the specialization of the governing equations to the axisymmetric coordinate is provided in the SOM along with a table of notation and parameter values (Tables S1, S2, and S3).

### 2.3 Numerical implementation

In the axisymmetric coordinates, the equations of motion (Eq. S14 and Eq. S15) simplify to a system of first-order differential equations (Eq. (S26)) with six prescribed boundary conditions (Eq. (S27)). In order to solve these coupled equations, we used the commercially available finite element solver COMSOL MULTIPHYSICS ^®^ 5.3. In this work, we assume that the total area of the membrane is conserved and to focus on the net effect of membrane tension, we set the transmembrane pressure at zero (*p* = 0).

## 3 Results

### 3.1 Beads formation along a nanotube due to protein induced spontaneous curvature

For membrane nanotubes, it has been shown that the composition of the lipid bilayer is a critical factor in determining their shapes and radii [69, 85, 29, 86]. To explore how a heterogeneity in the membrane properties due to a surface protein aggregation affects the shape of a nanotube, we conducted simulations on cylindrical nanotubes with a significant aspect ratio of radius 35 nm and length 100 *μ*m, and set the boundary tension to λ_0_ = 0.064 pN/nm. The effect of boundary tension on the initial nanotube radius and length is shown in Figs. S1 and S2. On this nanotube with homogeneous bending rigidity (*κ* = 320 pN.nm), we prescribed a surface distribution of the protein density. To have a smooth continuous transition, we implemented the difference in the protein density using a hyperbolic tangent function Eq. (S28), such that the area covered by the proteins at the center of the nanotube remains constant (A_protein_ = 1.6π*μ*m^2^), and the number of proteins per area increases from *σ*_0_ = 0 to *σ*_0_ = 1.25 × 10^−4^ nm^-2^ (Fig. 2A). As shown in Fig. 2B, the membrane bends outward in the area of the protein aggregation in response to the protein-induced spontaneous curvature. As the number of proteins in the defined area is increased, the membrane deformation also increases such that it resembles a bead-shaped structure form along the nanotube (Fig. 2C). The local variation of membrane tension corresponding to the membrane bending in the beaded domain and the applied incompressibility constraint is shown in Fig. S3.

**Fig. 2:**
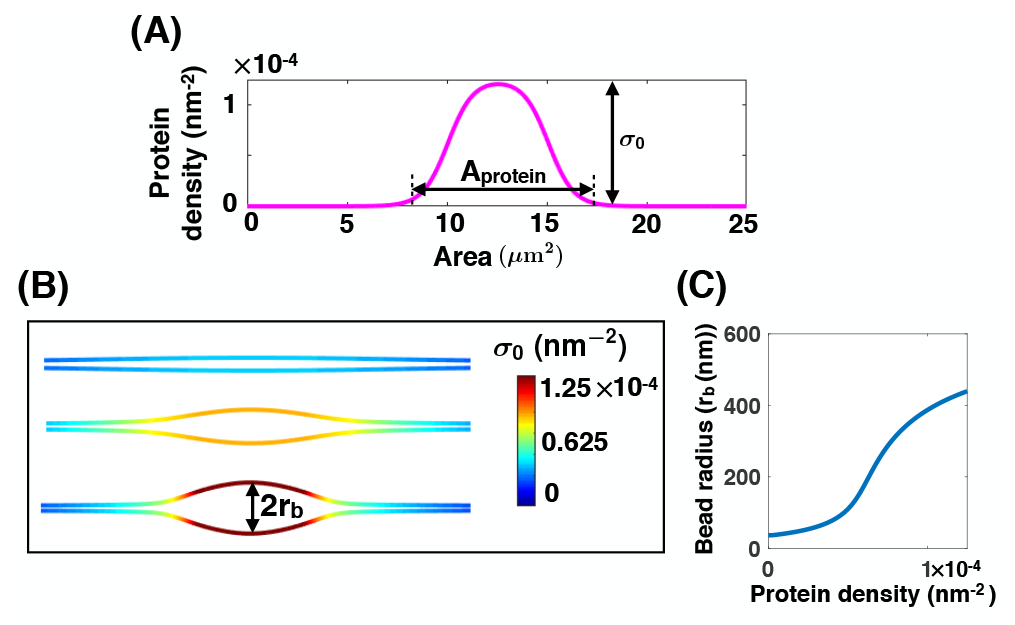
Protein-mediated bead formation along a membrane nanotube. (A) Protein density distribution on the membrane surface in which A_protein_ = 1.6π *μ*m^2^ shows the area coverage of the protein-enriched domain and *σ*_0_ represents the number of the proteins per unit area. (B) The formation of a large bead-shaped structure along the membrane nanotube as the density of proteins (*σ*_0_) increases for λ_0_ = 0.064pN/nm and uniform bending rigidity. (C) Bead radius increases as a function of the protein density.

### 3.2 Heterogeneity in membrane stiffness lead to the formation of bead-like structures along a nanotube

Motivated by our numerical observation that a protein induced spontaneous curvature along the membrane can result in the bead-like structures, we next asked if a change in the membrane stiffness due to membrane protein interaction could also induce a similar deformation along the nanotube. To answer this question, we repeated the simulation in Fig. 2 for *σ*_0_ = 1.25 × 10^−4^ nm^-2^ assuming that the bending rigidity along the covered area by the proteins is higher that the rest of the membrane (defining *κ*_ratio_ = *κ*_rigid_/*κ*_lipid_), but *C* = 0 (e.g for cylindrical proteins) [82, 86] (Fig. 3A). This represents a case where membrane-protein interaction induces a change in the membrane composition but does not induce an asymmetry between the leaflets. With increasing *κ*_ratio_ from *κ*_ratio_ = 1 to *κ*_ratio_=30 [82, 86, 87], the membrane bends significantly in the area of the rigid segment and an ellipsoidal bead-shaped structure forms along the nanotube (Fig. 3B). For *κ*_ratio_ < 10, the radius of the bead increases almost linearly. However, for *κ*_ratio_ > 10 the size of the bead remains almost constant (Fig. 3C). The local membrane tension distribution along the nanotubes and the bead-shaped structures is shown in Fig. S4. For completeness, we also applied a variation in the Gaussian modulus along the area of the protein aggregation (*Δκ_G_* = (*κ*_*G*,protein_ — *κ_G_*_,iipid_)/*κ_G_*_,iipid_). With varying —20 < *Δκ_G_* < 20, in the case that *σ*_0_ = 1.25 × 10^−4^ nm^-2^, *C* = 0, and *κ*_ratio_ = 1, we found that the effect of varying Gaussian modulus alone leads to insignificant membrane deformations (see Fig S5).

**Fig. 3:**
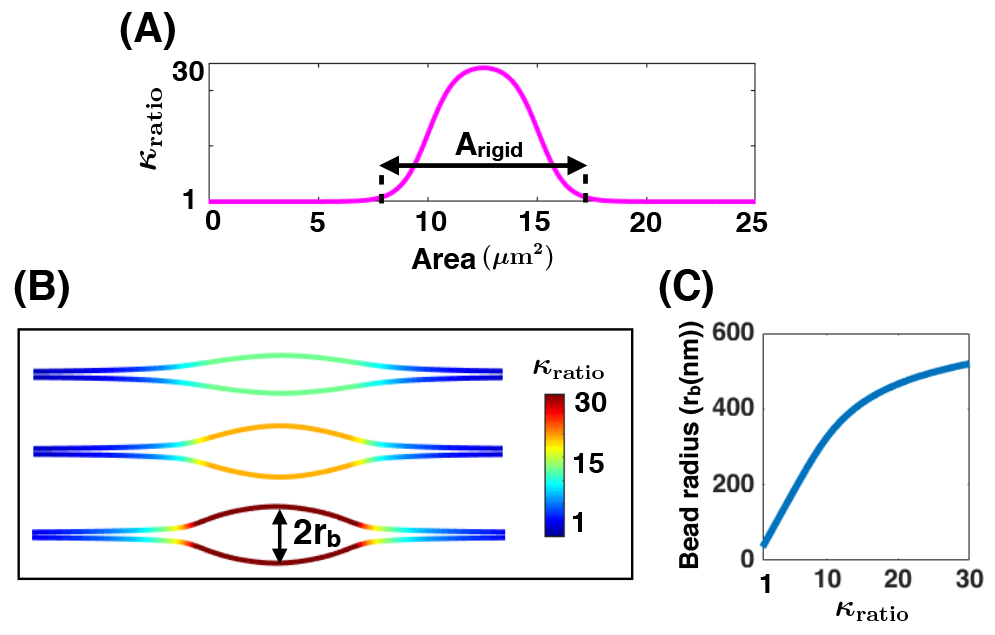
Heterogeneous membrane properties result in the formation of local bead-shaped structures (here σ_0_ = 1.25 × 10^−4^ nm^-2^). (A) Bending modulus variation along the area of the protein aggregation. *κ*_ratio_ shows the bending rigidity ratio of the rigid protein domain compared to that of the bare lipid membrane. (B) Membrane deformation in the region of large bending rigidity resembles a local bead formation phenomena; the tension at the boundary is set as λ_0_ = 0.064 pN/nm. (C) Bead radius increases as a function of *κ*_ratio_.

**Table 1:**
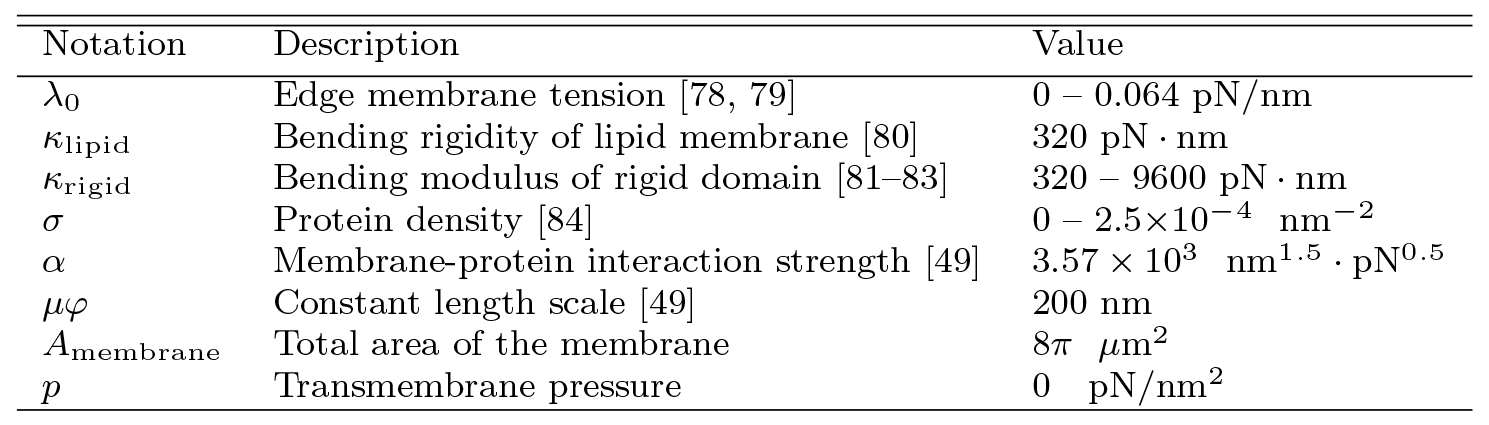
Parameters used in the model

| Notation | Description | Value |
| --- | --- | --- |
| $\lambda_0$ | Edge membrane tension [78, 79] | 0 – 0.064 pN/nm |
| $\kappa_{\text{lipid}}$ | Bending rigidity of lipid membrane [80] | 320 pN · nm |
| $\kappa_{\text{rigid}}$ | Bending modulus of rigid domain [81–83] | 320 – 9600 pN · nm |
| $\sigma$ | Protein density [84] | 0 – $2.5 \times 10^{-4}$ nm$^{-2}$ |
| $\alpha$ | Membrane-protein interaction strength [49] | $3.57 \times 10^3$ nm$^{1.5}$ · pN$^{0.5}$ |
| $\mu\varphi$ | Constant length scale [49] | 200 nm |
| $A_{\text{membrane}}$ | Total area of the membrane | $8\pi$ $\mu\text{m}^2$ |
| $p$ | Transmembrane pressure | 0 pN/nm$^2$ |

### 3.3 Morphological landscape of bead-shaped structures along a nanotube

Thus far, simulation results demonstrate that two unrelated mechanisms, protein-induced curvature, and heterogeneity in the membrane rigidity, independently lead to the formation of the bead-like structures along the membrane nanotube and that the radius of the bead increases nonlinearly with increasing strength of the heterogeneity. In order to explore how these two mechanisms might interact and modulate the shape of a nanotube, we conducted simulations where the heterogeneous domain has effects both from protein-induced spontaneous curvature and from increased bending rigidity. Initially, we repeated the simulations in Fig. 2 but this time assuming that the bending rigidity is heterogeneous (*κ*_ratio_ = 11). Interestingly, we found that the interaction of these two mechanisms leads to the formation of beads with different shapes. Based on the magnitude of the protein density, three different oblate spheroid shapes were obtained – (*i*) an ellipsoidal bead at *σ*_0_ = 10^−5^ nm^-2^, (*ii*) a flat cylindrical bead at *σ*_0_ = 4 × 10^−5^ nm^-2^, and (*iii*) a large unduloid-shaped bead at *σ*_0_ = 5.5 × 10^−5^ nm^-2^ (Fig. 4A). These three different shapes of beads are classified based on the derivative of the membrane curvature at the center of the beads. In the ellipsoidal bead, the derivative of the curvature is positive. In the cylindrical bead the derivative of the curvature is zero, and in the unduloid-shaped bead, the derivative of the curvature is negative. Observing these different shapes of beads shown that the coupling between two modes of spatial heterogeneity along a membrane nanotube not only increases the radius of the bead (see Fig. S6) but also changes the morphology of the bead. Furthermore, this shape transition suggests that the energy landscape of the membrane, which is now modulated by a heterogeneity in *κ* and a heterogeneity in *σ*, can lead to a range of shapes. To further identify over what range of protein density and *κ*_ratio_ these shape transitions occur, we performed the same simulation over a range of protein densities, (*σ*_0_ =0 — 2.5 × 10^−4^ nm^-2^), as well as over a range of *κ*_ratio_ = 1 — 11, encompassing soft protein domains to very stiff clusters. This variation allowed us to construct a phase diagram to identify the regions of different bead morphologies (Fig. 4B). The red region represents the formation of ellipsoidal bead-shaped structures, the blue region denotes the cylindrical beads, and the green region indicates the unduloid-shaped beads configuration.

**Fig. 4:**
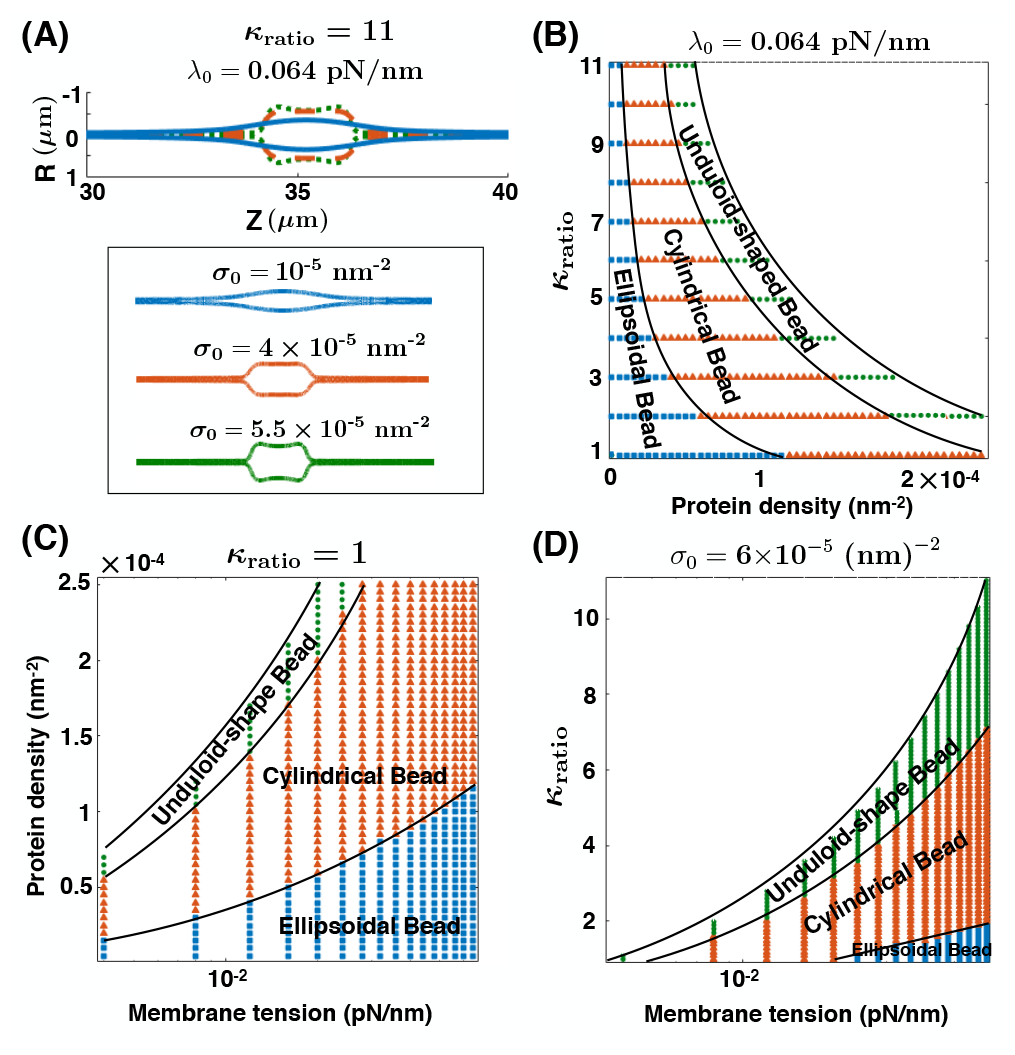
Bead morphology depends on the protein density (σo), the bending rigidity ratio of the protein-enriched domain compared to the lipid membrane (*κ*_ratio_), and the edge membrane tension λ_0_ (A) Three different possible shapes of a bead-like structure due to the presence of a rigid protein domain (*κ*_ratio_ = 11); (*i*) An ellipsoidal bead (red) at low protein density, (*ii*) a cylindrical bead (blue) at average protein density, and (*iii*) an unduloid-shaped bead (green) at high protein density. (B) Phase diagram for bending rigidity ratio versus the number of the proteins per unit area, λ_0_ = 0.064 pN/nm. The three different regions can be distinguished by the dominant length scale: (*i*) ellipsoidal beads when *L_σ_* ≫ *L_k_*, (*ii*) cylindrical beads when *L_σ_ ~ L_k_*, and (*iii*) unduloid-shaped beads when *L_k_* ≫ L_σ_ · (C) The protein density versus the edge membrane tension λ_0_ phase diagram for Kratio = 1· (D) The bending rigidity ratio versus the edge membrane tension λ_0_ phase diagram for *σ*_0_ = 6×10^−6^ nm^-2^. The colors in panels (C) and (D) represent the same bead shapes as panel (B).

These changes of the bead morphology can be better understood by comparing the two “induced” length scales by the rigid domain 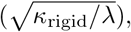 [42] and the protein aggregation (*L_σ_* = 1/*μφσ*) [34, 52]. The “induced” length scale by the membrane-protein interaction 1/*μφσ* is the preferred curvature of the protein-enriched domain and increases with decreasing *σ* (with *σ* = 0, both spontaneous curvature and the effect of protein aggregation are eliminated). The phase diagram in Fig. 4B shows that these two length scales act in tandem to regulate the bead size and shape. In order to compare the length scales, we consider a case with a fixed value of *L_k_* (e.g. *κ*_ratio_ = 2), and vary *L_σ_* by increasing the protein density (Fig. 4B). First, we see that the ellipsoidal beads form for low protein density, *i.e.* when *L_σ_* ≫ *L_k_* (Fig. 4B). Second, the cylindrical beads are energetically favorable for average values of protein density when the two length scales become comparable (*L_σ_* ~ *L_k_*). Finally, at very large values of protein density, when the “induced” length scale by the rigid domain becomes dominant (*L_k_* ≫ *L_σ_*), the unduloid-shaped bead forms along the nanotube (Fig. 4B).

Regardless of which mechanism dominates, the edge tension λ_0_ implicitly governs the length scale of the membrane; therefore the natural question that arises is how does this tension govern the shape and length scale transitions of the beads? We explore these questions by conducting two sets of simulations. First, for a homogeneous membrane (*κ*_ratio_ = 1), we varied the edge membrane tension (*λ*_0_) and the protein density (*σ*_0_) in a range (λ_0_ = 0.004 – 0.064 pN/nm and *σ*_0_ = 0 – 2.5 × 10^−4^ nm^-2^) (Fig. 4C). We observed that high edge tension shifted the transition of ellipsoidal to cylindrical and unduloid-shaped beads to the large protein densities. Second, we fixed the protein density (*σ*_0_ = 6 × 10^−5^ nm^-2^) and varied the edge tension (λ_0_) and the rigidity ratio (κ_m_tīo) between 0.004-0.064 pN/nm and 1-11 respectively (Fig. 4D). As our results show, in this case, all three possible shapes of beads are only accessible at high membrane tension. In general, we can see that by increasing the edge membrane tension either at a constant protein density or a fixed rigidity ratio, we decrease the value of the induced length scale by the rigid domain 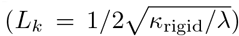, and therefore we move from the cylindrical and unduloid-shaped beads to the ellipsoidal bead-shaped region of the phase space.

### 3.4 Interaction between multiple beads along a nanotube

Often, multiple beads are observed along a membrane nanotube, suggesting that multiple domains of heterogeneity exist along the nanotube [15, 19, 73, 44-47, 17]. (Fig. 1A). These observations lead to the following question: how does the profile of these beaded strings depend on the different length scales associated with beaded nanotubes? In order to answer this question, we conducted simulations for the formation of two beads along the membrane by prescribing two areas of heterogeneity for three cases – (i) varying membrane rigidity alone in each domain in the absence of protein-induced spontaneous curvature, (ii) varying protein density for uniform rigidity, and (iii) varying protein density for domains with higher bending rigidity respect to the bare lipid membrane.

We first found when there are two regions of proteins far from each other with an end-to-end area given by *A*_separation_ = 1.6π *μ*m^2^ (Fig. 5A, top), two independent beads form (Fig. 5A bottom). The size and the shape of the beads are independent of the number of domains as long as the domains are far away from each other (Fig. S7 and Fig. S8). Having established that the regions of heterogeneity are independent when they are far from each other, we next asked under what conditions might these beads interact with one another? In other words, what length scales govern the stability features of multiple beads knowing that there is a certain relaxation length between the beads and cylinder? To answer this question, we repeated the simulation of two beads (Fig. 3), and varying the rigidity ratio (*κ*_ratio_) and end-to-end area (*A*_separation_) between 1-11 and 0 — 1.6π *μ*m^2^ respectively. Based on the results, we constructed a phase diagram separating the two possible morphologies; (*i*) two distinct beads represented by the color blue, and (*ii*) one single bead denoted by color red (Fig. 5B). We found that when the two domains are bought together by decreasing the area between the heterogeneity, there is a smooth transition from two beads connected by a string to a single bead (Fig. 5B, C). This suggests that at close distances, the energy minimum of the nanotube is attained for a single large bead rather than for two beads connected by a string. We also observe the smooth transition in the number of beads is accompanied by the shape transition from an unduloid-shaped bead to a large ellipsoidal bead (Fig. 5C).

**Fig. 5:**
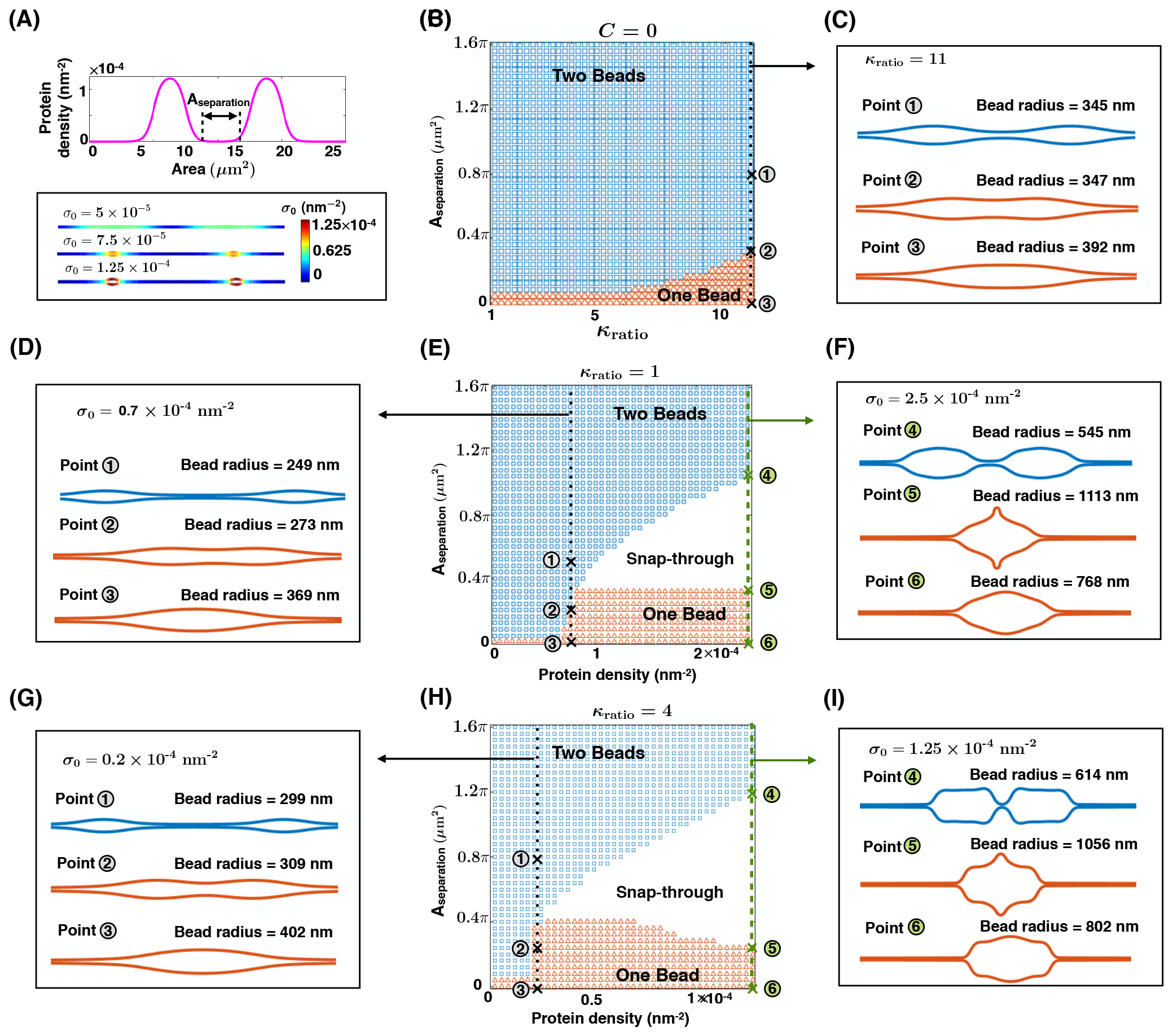
Multiple beads along a nanotube. (A, top) Protein density distribution with two domains of protein accumulation. The area of each protein-enriched domain is *A*_protein_ = 1.6π *μ*m^2^ and the domains are far from each other *A*_separation_ = 1.6π *μ*m^2^. The number of proteins within each domain increases by the same amount. (A, bottom) Two beads form corresponding to each protein-enriched domain, λ_0_ = 0.064 pN/nm. (B) The area separation between two beads versus the bending rigidity ratio phase diagram with *C* = 0. There are two possible shapes, (*i*) two separated beads denoted with the color blue, and (*ii*) one single bead marked by the color red. The transition from two to one bead is smooth everywhere in this parameter space. (C) The smooth membrane shape transition from two beads to one bead with decreasing area separation for *κ*_ratio_ = 11 and membrane profiles for points marked in panel (B). (D) Membrane profiles for the black dotted line in panel (E) show the smooth evolution of membrane shape from two beads to one bead at *σ*_0_ = 0.7 × 10^−4^ nm^-2^ with decreasing *A*_separation_ (*κ*_ratio_ = 1). (E) Phase diagram for the area separation between two beads versus the protein density at *κ*_ratio_ = 1. The colors represent the same morphologies as panel (B). Smooth transition from two beads to one bead at low protein density and a snap-through instability in the transition from two beads to one bead for large protein densities is observed. (F) Membrane profiles show the snap-through transition from two kissing beads to one large bead at *σ*_0_ = 2.5 × 10^−4^ nm^-2^ for the green dashed line in panel (E). (G) Membrane profiles for the black dotted line in panel (H) show the smooth evolution of membrane shape from two beads to one bead at *σ*_0_ = 0.2 × 10^−4^ nm^-2^ for *κ*_ratio_ = 4 with decreasing *A*_separation_. (H) Phase diagram for the area separation between two beads versus the protein density at κ_ratio_ = 4. The colors represent the same morphologies as panel (B). Smooth transition from two beads to one bead at low protein density and a snap-through instability in the transition from two beads to one bead for large protein densities are observed. (I) Membrane profiles show the snap-through transition from two kissing beads to one large bead at *σ*_0_ = 1.25 × 10^−4^ nm^-2^ for the green dashed line in panel (H).

However, when we varied the area separation between the beads for a range of protein densities in Fig.5A, we found a sharp transition from two beads to one bead accompanied by a snap-through instability (Fig. 5D, E, and F). For small protein densities, when the area between the two beads is decreased, the nanotube appears to have one large bead, while the transition from two beads to one bead is continuous indicating that there is no energy barrier to move from one state to another (Fig. 5D and Fig. 5E, black dotted line). However, as the protein density increases, we found the emergence of a snap-through instability for small separation areas (Fig. 5E) corresponding to bead shapes that show a distinct transition from two ellipsoids to a flower-shaped bead to a large ellipsoidal bead (Fig. 5E, green dashed line and Fig. 5F). This means that the landscape between two beads and one bead at high protein densities is governed by a large energy barrier (Fig. S9). And finally, when we repeated these calculations for a rigid protein domain such as *κ*_ratio_ = 4 (Fig. 5 G, H, I), we found that the area and protein density still govern the energy and stability landscape but the transition point – where the snap-through instability occurs – is shifted towards the lower protein densities, with no change in bead shapes (Fig. 5G, H, I). Thus, individual beads along nanotubes potentially tend to be dominant because any fusion event of two beads may often be accompanied by energy barriers and non-trivial shape transitions. This energy barrier corresponds to the competition between the length scale of the domain separation and the induced length scale due to the heterogeneity (see Fig. S10).

## 4 Discussion

Tunneling nanotubes are membranous projections between cells [1, 2, 17]. Much of the biophysics associated with these dynamic structures are only now beginning to be explored but it is becoming increasingly clear that the cellular membrane and membrane-protein interactions play a critical role in maintaining these cellular architectures [15, 69, 19, 85, 88]. In this study, we explored how the energy landscape and the role of heterogeneity in the membrane either due to protein aggregation or material properties alter the architecture of nanotubes. Our results can be summarized as follows – membrane heterogeneity due to either protein-induced spontaneous curvature (Fig. 2) or membrane rigidity (Fig. 3) can result in the formation of bead-like structures along a nanotube. Additionally, the interaction between these two modes of heterogeneity can lead to the formation of beads with distinct shapes while the transitions between these shapes from ellipsoidal to cylindrical to unduloid-shaped beads are consequences of competing length scales in the system (Fig. 4). Finally, the length scale competition induced by these effects coupled with the length scale between two regions of heterogeneity also governs the stability landscape of multiple beads (Fig. 5). Although a similar interplay between adhesion and bending energy has been explored in shaping the plasma membrane, [89, 90], the shaping of membrane nanotubes resulting from the interplay between a spatially varying membrane rigidity and protein-induced spontaneous curvature had not been previously been explored.

The change in the morphology of the membrane nanotubes from cylindrical tubules to the beaded architectures due to local variation in spontaneous curvature or membrane properties (Fig. 2 and Fig. 3) is reminiscent of the classic Rayleigh-Plateau instability [22-24]. In Rayleigh-Plateau instabilities, surface tension is the key parameter that regulates the growth rate of the sinusoidal perturbations along a column of a fluid [22-24]. Previous studies demonstrated that inducing a constant homogeneous membrane tension all along a cylindrical membrane can also lead to a dramatic shape transformation into a modulated structure of a string of pearls [25-30]. However, the tension of lipid membranes can change not only globally but also locally due to the absorption of proteins, nanoparticles, inclusions, or actomyosin interactions with the membrane [34-38]. Here, we show that local variation of the membrane tension corresponding to the membrane heterogeneities in the beaded nanotubes (Figs. S3 and S4) may play a role in governing the morphology of these tubes.

Our simulations allow us to make the following predictions. Tension at the edge of the nanotube not only governs the nanotube radius but also its response to heterogeneity. Therefore, manipulating cell tension and evaluating how it affects the morphology of the nanotube will provide information on how the tensional homeostasis of cells affects the membrane nanotubes. Additionally, we found that there is an energy barrier that governs the landscape of the transition from two beads to one bead. This energy barrier, governed by a snap-through instability suggests that the fusion of two beads depends on the membrane composition and its material properties such that under high protein density or high rigidity conditions, there is a large crossover energy for fusion. These predictions can be extended to multiple beads as well.

In the broader context of interactions between the bilayer membrane and curvature inducing moieties (proteins and cytoskeletal filaments), Shlomovitz and Gov showed that the coupling between membrane shapes and membrane-bound FtsZ filaments can induce high-density FtsZ rings along a cylindrical membrane [73]. With no entropic effects, these rings interact with each other, can coalesce and form larger rings depending on the membrane tension and separation distances. They predict that when the separation between two rings in large than 2πR (R is the radius of the cylinder), membrane shape undulations around each ring act as an energy barrier to stabilize the separate rings and preventing coalescence [73]. These results suggest that the observed energy barriers in our model and Shlomovitz and Gov paper could have the same origin since both responses appear due the elastic behavior of the lipid membranes in interactions with local curvature inducing moieties. Additionally, recent experiments on the fission of yeast have demonstrated that the formed rings along the tubular membranes by the actin-myosin contractile force interact and fuse when the natural width of the ring is much smaller than their separation distance [73, 91]. This is also consistent with our simulation results, where we find that the transition from two beads to one bead occurs when the distance between two beads is one order of magnitude smaller than the induced length scale by the heterogeneity (Fig. S10).

Although our model has provided some fundamental insights into the role of membrane composition in nanotube morphology, we have made simplifying assumptions that will need to be revisited as more experimental evidence is gathered regarding these structures. Additionally, the dynamics of membrane-protein diffusion and what could be the underlying mechanisms that govern protein aggregation along the nanotube as suggested in our model are not yet fully understood. While there is evidence for the strong curvature-mediated feedback for the protein aggregation [63], it is possible that feedback between proteins in the lumen of the nanotube and biochemical signaling can lead to the formation of protein microdomains. For example, it is known that phase separation between two main components of the membrane – clusters of sphingolipids and cholesterol molecules – can result in the formation of lipid rafts [82, 92]. In addition, the role of the cytoskeleton (actin and microtubule filaments) and motor protein transport along the nanotube is known to be important [1, 2, 93], but a correlation with the beading morphology is yet to be established. Also, there are only a handful of direct experimental observations of membrane heterogeneities along nanotubes in cells [44]. While it is easy to imagine the beaded morphologies in *in vitro* studies [20, 45], it remains to be seen if such morphologies will be observed in cells. And finally, the role of the active transport versus the flow of cytosolic components in governing the stability of these nanotubes is not captured by our model and remains an active focus of our research and modeling efforts.

## Acknowledgment

This work was supported by ARO W911NF-16-1-0411 grants to P.R. H.A. was supported by a fellowship from the Visible Molecular Cell Consortium (VMCC), a program between UCSD and the Scripps Research Institute. B.O. was supported by NIH R01GM111938. The authors would also like to thank Prof. Sasa Svetina and Dr. Morgan Chabanon for their comments and help on improving the manuscript.

## Author contributions

B.O. and P.R. conceived the research, H.A. and P.R. conducted the research and analyzed the data, H.A., B.O., and P.R. wrote the paper. All authors reviewed the manuscript and agreed on the contents of the paper.

